# A double-hit *in vivo* model of *GBA1* viral microRNA-mediated downregulation and human alpha-synuclein overexpression demonstrates nigrostriatal degeneration

**DOI:** 10.1101/2021.06.23.449545

**Authors:** Alexia Polissidis, Georgia Nikolopoulou, Effrosyni Koronaiou, Maria Nikatou, Catherine Viel, Marios Boyongo, S. Pablo Sardi, Maria Xilouri, Kostas Vekrellis, Leonidas Stefanis

**Author notes:** 2^nd^ Neurological Department, Henry Dunant Hospital Center, 107 Mesogeion Ave, 11526, Athens, Greece (Georgia Nikolopoulou). To whom correspondence should be addressed: Alexia Polissidis, Leonidas Stefanis, Center of Clinical, Experimental Surgery and Translational Research, Biomedical Research Foundation of the Academy of Athens (BRFAA), Soranou Efesiou 4, Athens, 11527, Greece.

## Abstract

Preclinical and clinical studies support a strong association between mutations in the *GBA1* gene that encodes β-glucocerebrosidase (GCase) (EC 3.2.1.45; glucosylceramidase beta) and Parkinson’s disease (PD). Alpha-synuclein (AS), a key player in PD pathogenesis, and *GBA1* mutations may independently and synergistically cause lysosomal dysfunction and thus, embody clinically well-validated targets of the neurodegenerative disease process in PD. However, double-hit *in vivo* models, recapitulating pathological features of PD that can be used to dissect the nature of the complex relationship between GCase and AS on the nigrostriatal axis, the region particularly vulnerable in PD, are direly needed. To address this, we implemented a bidirectional approach in mice to examine the effects of: 1) GCase overexpression (wildtype and mutant N370S) on endogenous AS levels and 2) downregulation of endogenous GCase combined with AS overexpression. Striatal delivery of viral-mediated GCase overexpression revealed minimal effects on cortical and nigrostriatal AS tissue levels and no significant effect on dopaminergic system integrity. On the other hand, microRNA (miR)-mediated *GBA1* downregulation (miR GBA), combined with virus-mediated human AS overexpression (+AS), yields decreased GCase activity in the cortex, mimicking levels seen in *GBA1* heterozygous carriers (30-40%), increased astrogliosis and microgliosis, decreased striatal dopamine levels (50% compared to controls) and loss of nigral dopaminergic neurons (~33%)-effects that were all reversible with miR rescue. Most importantly, the synergistic neurodegeneration of miR GBA+AS correlated with augmented AS accumulation and extracellular release in the striatum. Collectively, our results suggest that GCase downregulation alone is not sufficient to recapitulate key pathological features of PD *in vivo*, but its synergistic interplay with AS, via increased AS levels and release, drives nigrostriatal neurodegeneration. Furthermore, we demonstrate a novel double-hit model that can be used to identify putative mechanisms driving PD pathophysiology and can be subsequently used to test novel therapeutic approaches.

## Introduction

*GBA1* gene mutation (hereon referred to as GBA) heterozygosity is one of the most common genetic risk factors for Parkinson’s disease (PD) with a predicted frequency of 7% to 10% in the PD population (Neumann et al., 2009; Schapira, 2015; Sidransky et al., 2009). The GBA gene encodes for glucocerebrosidase (GCase), a lysosomal enzyme responsible for the breakdown of glucosylceramide (GlcCer) to glucose and ceramide. GBA-associated PD (GBA-PD) exhibits a comparable symptomatic presentation to sporadic PD but with more rapid cognitive and motor decline and a slightly earlier average age of onset (Alcalay et al., 2012; Brockmann et al., 2015; Oeda et al., 2015). Several clinical studies provide compelling evidence for a reciprocal detrimental relationship between GCase (loss of function) and aberrant alpha-synuclein (AS) (toxic gain of function). For example, heterozygous mutant *GBA1* carriers demonstrate increased post-mortem levels of AS, while sporadic cases of PD have decreased brain GCase activity associated with early accumulation of AS, inflammation, dysregulation of chaperone-mediated autophagy and lipid metabolism (Chiasserini et al., 2015; Gegg and Schapira, 2018; Murphy et al., 2014).

The central role of AS accumulation in GBA-PD is emphasized by the reciprocal relationship between GCase activity and AS demonstrated in SH-SY5Y cell cultures, neuronal cultures, conduritol-b-epoxide (CBE)-treated mice and transgenic GBA mouse models (Cullen et al., 2011; Manning-Boğ et al., 2009; Mazzulli et al., 2011; Migdalska-Richards et al., 2017; Osellame et al., 2013; Sardi et al., 2013; Sidransky and Lopez, 2012, 2012; Tayebi et al., 2017; Xu et al., 2011). In addition, several preclinical studies have demonstrated the successful amelioration of neurodegenerative phenotypes in murine synucleinopathy models by either enhancing GCase activity in the central nervous system (Sardi et al., 2013) or by targeting the GCase pathway indirectly via inhibition of glucosylceramide synthase (Sardi et al., 2017). Conversely, cell and mouse models have demonstrated that increased AS causes a decrease in GCase activity (Gegg et al., 2012; Migdalska-Richards et al., 2017).

Considering the prevalence of *GBA1* mutations and the role both GCase and AS play in PD, understanding how GBA affects the susceptibility to AS-mediated nigral toxicity *in vivo* is crucial for understanding PD pathogenesis and successfully slowing or blocking the underlying neurodegenerative process with future disease-modifying treatments. However, the overall utility of currently available murine models is limited (reviewed extensively in Farfel-Becker et al., 2019). Here, we sought to recapitulate key pathological PD features, i.e. nigral neurodegeneration, neuroinflammation, and AS accumulation, to generate a model of GBA heterozygosity that more closely represents the haploinsufficiency as seen in GBA carriers and assess its association with AS accumulation and PD pathogenesis.

## Materials and Methods

### *In vitro* experiments

For the *in vitro* experiments, HEK293T cells were double transfected with AAV plasmids carrying the human GBA (wild-type (WT) or mutant N370S, D409V, L444P) or GFP (as control group) together with human AS using the calcium phosphate method in separate concentrations of 0.75 μg (1:1 ratio). After transfection, the plate was incubated at 39° C for 24 hours in 1-% FBS DMEM (plasmids contain a Gus B thermosensitive promoter). The plate was then incubated at 37° C for 48 hours in 2% FBS DMEM and the medium was switched from full to 2% FBS. The medium was collected and spun down twice at 2000 and 4000 rpm and the cells were lysed in Triton-X soluble buffer and further analyzed by Western Blot analysis and ELISA.

For the *in vitro* experiments with microRNA GBA plasmid, Neuro2A cells were transfected with lipofectamine 2000 (Invitrogen) following the manufacturer’s instruction vs. GFP as control (Table 1). The cells were incubated as above for HEK293T cells. The cells were collected 72 h post-transfection and lysed in Triton-X soluble buffer and analyzed by Western Blot.

**Table 1:** Names and titers of AAVs injected in C57Bl/6 mice striata; miR= microRNA, AS= α-synuclein.

| Treatment (abbreviated) | Virus name | Virus Titer (viral particles (vp)/μl) |
| --- | --- | --- |
| **Experiment 1: GBA overexpression** |  |  |
| GFP | rAd-GFP | $1.51 \times 10^8$ |
| wt GBA | rAd-WT GBA | $2.31 \times 10^8$ |
| N370S | rAd-N370S | $5.9 \times 10^8$ |
| **Experiment 2: GBA downregulation & AS overexpression** |  |  |
| miR control | AAV2/1 CBA-miR-Control | $2.80 \times 10^{12}$ |
| miR control-GFP | AAV1 sc-GFP | $5.8 \times 10^{12}$ |
| miR GBA | AAV2/1 EGFP-miRNA-gba1 B | $3.30 \times 10^{13}$ |
| miR rescue | AAV2/1 scGusB-GC*-miR gbaB | $3.40 \times 10^{12}$ |
| AS | AAV1-CBA-aSynuclein | $6.30 \times 10^{12}$ |

### Animals

Eight-week-old male wild-type C57Bl/6 mice (27-33 g body weight) were housed with free access to food and water under a 12-h light/dark cycle. All experimental procedures performed were approved by the Ethical Committee for Use of Laboratory Animals of the Biomedical Research Foundation Academy of Athens and in accordance with the ARRIVE guidelines the EU Directive 2010/63/EU for animal experiments.

### AAV vectors

For experiment 1, recombinant human serotype 5 adenoviruses (rAd) expressing fulllength human wild-type (WT), D409V, L444P and N370S GBA were generated, as previously described (Papadopoulos et al. 2018, He et al. 1998). For experiment 2, adeno-associated viral vectors were used to downregulate GCase expression with an artificial microRNA (miR)-based hairpin targeting mouse *GBA1* (miR GBA), control vector with a scrambled miR (miR control) or co-transduced with GFP where indicated (miR control-GFP), rescue vector expressing a miR-resistant GBA (i.e. a series of synonymous mutations were introduced to the human GBA coding sequence to generate AAV2/1-GusB-GBA*-miR-Gba, where GBA* is resistant to the artificial miR) and vector overexpressing human AS have been previously described (Sardi et al., 2011, Jackson et al. 2019). For experiment 1, subjects were assigned to one of three treatment groups: GFP, WT and N370S; and for experiment 2, subjects were assigned to one of six treatment groups: miR control +/- AS, miR GBA +/- AS and miR rescue GBA +/- AS (see table 1 for the list of viruses and titers).

**Table 2:** Experiment 2 *in vivo* GBA downregulation: The 6 virus-injected experimental groups and their abbreviations; AS= α-synuclein.

| Treatment | Abbreviation |
| --- | --- |
| microRNA control (GFP) – human AS | miR CTL – AS |
| microRNA GBA (GFP) – human AS | miR GBA – AS |
| microRNA rescue – human AS | miR Resc – AS |
| microRNA control (GFP) + human AS | miR CTL + AS |
| microRNA GBA (GFP) + human AS | miR GBA + AS |
| microRNA rescue + human AS | miR Resc + AS |

### Surgical Procedures

All surgical procedures were performed under isoflurane (Abbott, B506) anesthesia. Animals were given i.p. carpofen (dose) for analgesia. After placing the animal into a stereotaxic frame (Kopf Instruments, USA), 2 μl of AAV solution (with final titer of 3.3 E13 gc/mL) and 1 μl of DPBS or human AS was injected unilaterally into the right striatum using the following coordinates: −0.5 mm anteroposterior, +1.6 mm mediolateral from the bregma, and −3.4 mm dorsoventral from the skull, according to the mouse stereotaxic atlas (Paxinos and Watson, 2006). Injections were performed using a finely pulled glass capillary (diameter of approximately 60–80 μm) attached to a Hamilton syringe with a 22 gauge needle. After delivery of the viral vector using an injection rate of 0.1 μL/15 sec, the capillary was held in place for 5 min, retracted 0.1 μm, and, after 2 min, was slowly withdrawn from the brain, as previously described (Polissidis et al., 2020).

### Immunohistochemistry

Four and eight weeks post-injection (WPI), animals were perfused under isoflurane anesthesia. The perfusion was performed through the ascending aorta using approximately 30 mL of 1X PBS for blood clearance and then 50 mL of 4% paraformaldehyde (PFA) for tissue fixation. The brains were removed and post-fixated overnight in 4% PFA, then transferred to 15% sucrose overnight followed by 30% sucrose overnight. The brains were then frozen using isopentane on ice (−55°C) and stored at −80°C until analysis. The perfused brains were cryosectioned through the coronal plane in 30 μm increments.

For immunohistochemical analysis, striatal and nigral sections were washed with PBS, followed by antigen retrieval with 10 mM citrate buffer at 80°C for 20 minutes, then placed on ice for an additional 20 minutes. The sections were then blocked using 5% normal goat serum (NGS) and 0.1% Triton-X for 1 hour at room temperature. Subsequently, the sections were incubated for 48 hours at 4°C with the following primary antibodies: anti-GBA (1:500; human 8E4; human and rodent G4171, Sigma-Aldrich), anti-human α-synuclein (syn211) (1:150,000;, 36-008, Millipore), anti-GFP (1:2000; 13970, Abcam), anti-TH (1:2000; AB152 MAB318, Millipore), anti-dopamine transporter (DAT; 1:2000; AB2231, Sigma-Aldrich), anti-GFAP (1:750, Z0334, DAKO) anti-AIF-1/Iba1 (1:1000, 019741 DAKO); and secondary antibodies: CF488A (1:2000; 20010, 20012 Biotium), CF555 (1:2000, 20030, 20033 Biotium), Affinipure Cy5 (1:400, 115-175-146 Jackson Immunoresearch), and DAPI (1:2000; Sigma).

### Total RNA isolation and reverse transcription

Midbrain tissue was dissected from mouse brains and total RNA was isolated. For this purpose, 1 mL of TRIZOL was used per 50-100 mg of tissue, followed by centrifugation at 12,000xg for 10 minutes at 4°C. The supernatant was transferred to a new tube and incubated for 5 minutes at room temperature. Afterwards, 0.2 mL of chloroform was added per 1 mL of TRIZOL reagent. The tubes were shaken vigorously for 15 seconds and incubated again at room temperature for 3 minutes, followed by centrifugation at 12000xg for 15 minutes at 4°C. The mixture was separated into a lower red, phenol-chloroform phase, an interphase, and a colorless upper aqueous phase, which contains the RNA. The aqueous phase was transferred to a new tube and the RNA was precipitated using 0.5 mL of isopropyl alcohol per 1 mL of TRIZOL. Samples were incubated overnight at −20°C and were centrifuged again at 12000xg for 10 minutes at 4°C and the supernatant was then completely removed, leaving only the RNA pellet which was washed using 75% ethanol (1:1 TRIZOL). After another centrifugation at 7500xg for 5 minutes at 4°C, the pellets were air dried and dissolved in RNase-free H_2_O. Concentration was measured using spectrometry and RNA integrity was assessed by running 500 ng of each sample on a 1% agarose gel.

DNase treatment was then performed (RQ1 10x buffer, RQ1 DNase by Promega, 30 minutes at 37°C) and stopped by adding 1 μL of DNase Stop Solution (Promega) to the sample and incubating for 10 minutes at 65°C. Finally, 1 μL of oligo dT per sample was added and the samples were incubated for 5 minutes at 70°C. For the cDNA synthesis, we used 5 μL of 5X RT buffer (Promega), 5 μL of dNTPs, 1 μL of RNaseOUT (Promega) and 1 μL of reverse transcriptase (M-MLV RT, Promega) and incubated the samples for 1 hour at 37°C. qPCR was then performed.

### Confocal microscopy and analysis

Images from the stained sections were obtained using confocal microscopy (Leica SP5 mark II with conventional photon-multiplier tube, at 23° C using the Leica Advanced Fluorescence v2.7 acquisition software (Leica Microsystems, Wetzlar, Germany)). After acquiring representative images, quantification of the stained proteins was done using Imaris (Version 8.0) for GBA or ImageJ for AS. For each animal, two sections per brain area were used and a total of 3 images per section were obtained.

For GBA quantification (n=5 for 4 WPI, n=3 for 8 WPI), the surface function of the Imaris software was used to select only GFP-positive cells by appropriately adjusting the number of voxels as well as the intensity of the GFP channel in the threshold tab. The intensity of GBA was then calculated by masking the GBA channel within the selected surfaces.

For the quantification of human AS (n=4-6) in the striatum and substantia nigra, the entire surface using DAPI intensity or TH positive somata, respectively, was selected and the threshold was adjusted appropriately, after converting the image to 8-bit. The selected area was assigned as the region of interest (ROI) and using the measure tool, intensity of human AS staining within the ROI was determined.

### Glucocerebrosidase activity measurements

Brain glucocerebrosidase activities were determined as previously described using 4-methylumbelliferyl-β-d-glucopyranoside (4-MU) as the artificial substrate in striatal tissue (Sardi et al., 2013) and cortical tissue (Papadopoulos et al., 2018).

### Glycosphingolipid level measurements

Striatal tissue levels of glucosylceramide (GlcCer), glucosylsphingosine (GlcSph) and ceramide (Cer) were measured by liquid chromatography and tandem mass spectrometry (LC-MS/MS) as previously described (Sardi et al., 2013).

### Behavioral analysis: cylinder test

Spontaneous locomotor activity was assessed at 4 and 8 WPI. Briefly, animals were placed in a Plexiglas cylinder (dimensions) and allowed to move freely for 5 min. Videos were recorded from below with a mobile phone camera (selfie mode) and total number of rearings (left, right, and both forelimb placements) and distance travelled (cm) were analyzed manually and with Ethovision XT9.0 software (Noldus).

## Results

### Experiment 1: GCase overexpression and measurement of endogenous AS levels

#### Mutant N370S GCase decreases secreted AS *in vitro*

Wild-type GBA (WT GBA) GCase overexpressing HEK293 cells demonstrated enhanced GCase enzymatic activity levels compared to the control group (p<0.0001) while the overexpression of the PD-linked mutant GCase forms (N370S, D409V, L444P) did not alter GCase enzyme activity (SF1A). Furthermore, intracellular AS levels remained largely unaffected by WT or mutant GCase overexpression (SF1B); however, mutant N370S GBA exclusively decreased the % of secreted (extra/intracellular) human AS (SF1C).

#### Mutant N370S and WT GCase overexpression affect neither nigrostriatal dopaminergic system integrity nor AS expression *in vivo*

Unilateral striatal injections of GFP, WT GBA or N370S AAVs in mice led to efficient delivery of transgenes to the striatum, expression in the cortex, and retrograde transport to the substantia nigra at 8 WPI (SF2). WT GBA overexpression led to a nonsignificant increase in net cortical GCase enzyme activity (Fig 1D) but no changes in striatal dopamine levels (Fig 1E) or tyrosine hydroxylase (TH) density of dopaminergic terminals (Fig 1F). Measurements of total AS protein levels in the striatal, cortical, and ventral midbrain tissue did not reveal any statistically significant changes following WT GBA or N370S GBA overexpression (Fig 1G-I).

**Figure 1.**
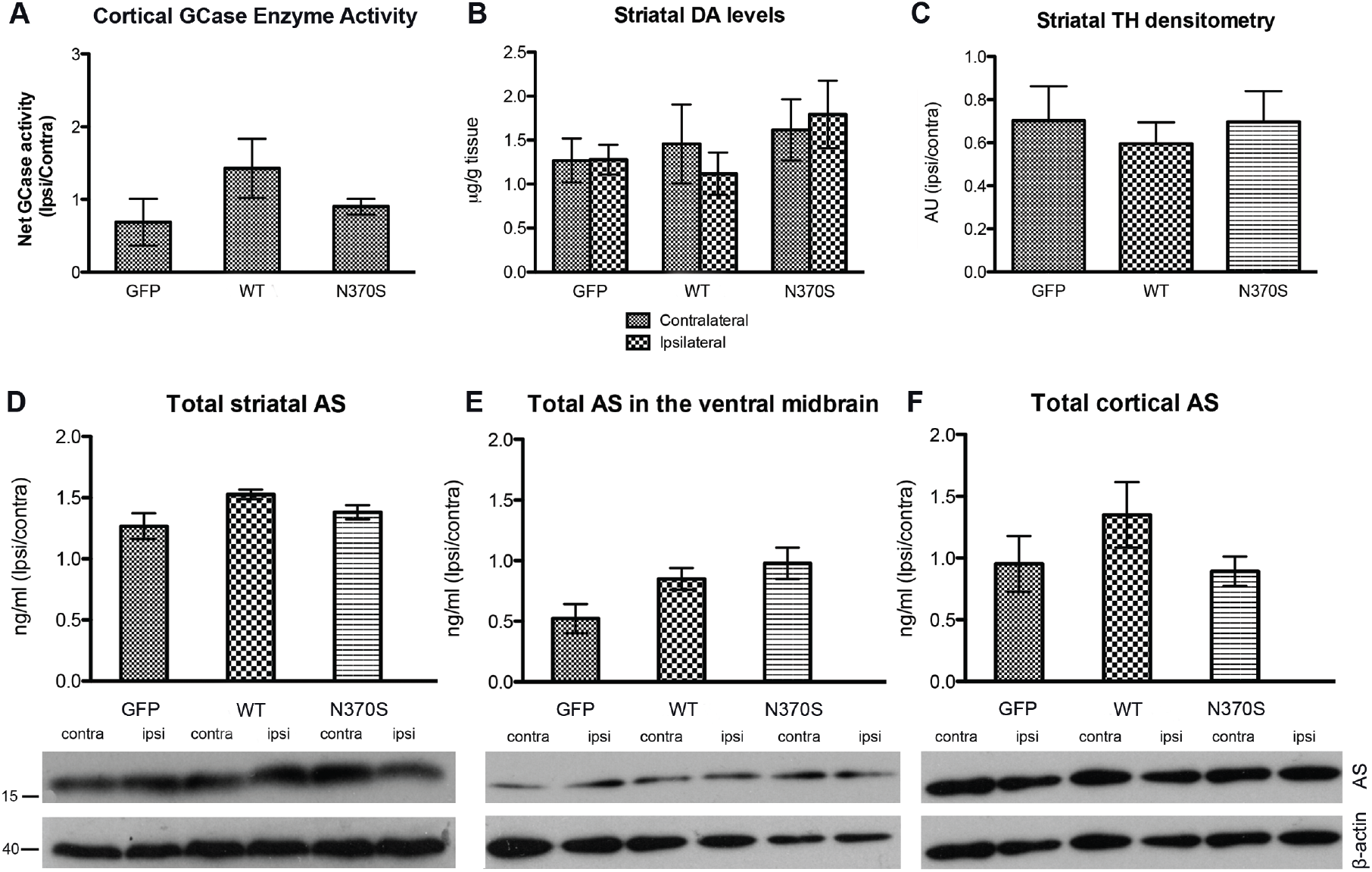
*In vivo* viral overexpression of GFP, wild-type (WT) and mutant (N370S) GBA. *In vivo* unilateral striatal overexpression of WT GBA or N370S GBA (vs. GFP) did not affect cortical GCase enzyme ipsilateral/contralateral activity (ipsi(lateral)/contra(lateral) activity); although there was a definite trend for an increase with WT GBA (A), striatal dopamine (DA) levels (expressed as μg/g wet tissue) (B), striatal tyrosine hydroxylase (TH) intensity (measured in arbitrary units (au) ipsi/contra) (C), total AS levels (ipsi/contra) in the striatum (D), ventral midbrain (E), and cortex (F) as measured by Western immunoblotting (top panels: quantification of AS (measured with the Syn1 antibody and normalized to β-actin levels, expressed ipsi/contra; bottom panels: immunoblots of AS (at 17 kDa) and β-actin (40 kDa)).

### Experiment 2: GCase downregulation and human AS overexpression

The *in vitro* experiments in HEK293T cells showed that GCase enzymatic activity was increased two fold in WT GBA-overexpressing cells (p<0.0001), while the three mutations showed similar GCase activity levels as GFP-expressing cells (SF1A). The D409V and L444P GBA mutations showed no significant changes to intracellular or extracellular AS levels, whereas a statistically significant reduction in secretion of AS in the presence of N370S mutated plasmid of GBA was observed without a significant change in intracellular levels of AS (SF1B,C). Therefore, the N370S mutation was chosen for the *in vivo* experiments.

#### Significant GCase downregulation at 8 WPI

miR GBA-GFP and human AS expressing AAVs infected the striatum, the site of injection, as well as the cortex and were efficiently retrogradely transported to the ipsilateral substantia nigra (Fig 2A,B). miR GBA-induced GCase downregulation was determined in Neuro2A cells (Fig 2C), and then *in vivo* at 4 and 8 WPI. No significant GCase downregulation was observed at 4 WPI in miR GBA-GFP-injected mouse striatum (SF3). However, GCase levels decreased by approximately 40% at 8 WPI, as assessed by immunohistochemistry in the ipsilateral striatum (t_4_=3.046, p=0.0382) (Fig 4A, D). A similar magnitude of cortical GCase activity reduction was observed in miR GBA-GFP injected mice (miR treatment effect: [F(_2,25_)= 9.42, p=0.0009], Fig 2E) and glucosylceramide accumulation was observed in the substantia nigra, which was reversed in miR Resc-injected mice (miR treatment effect [F(_2,53_)=25.13, p<0.0001], Fig 2F). Levels of other glycosphingolipids including sphingomyelin and psychosine were unchanged (SF4) and glycosylsphingosine levels were undetectable. Based on these results, further analysis of dopaminergic system integrity and AS pathology was performed at 8 WPI.

**Figure 2.**
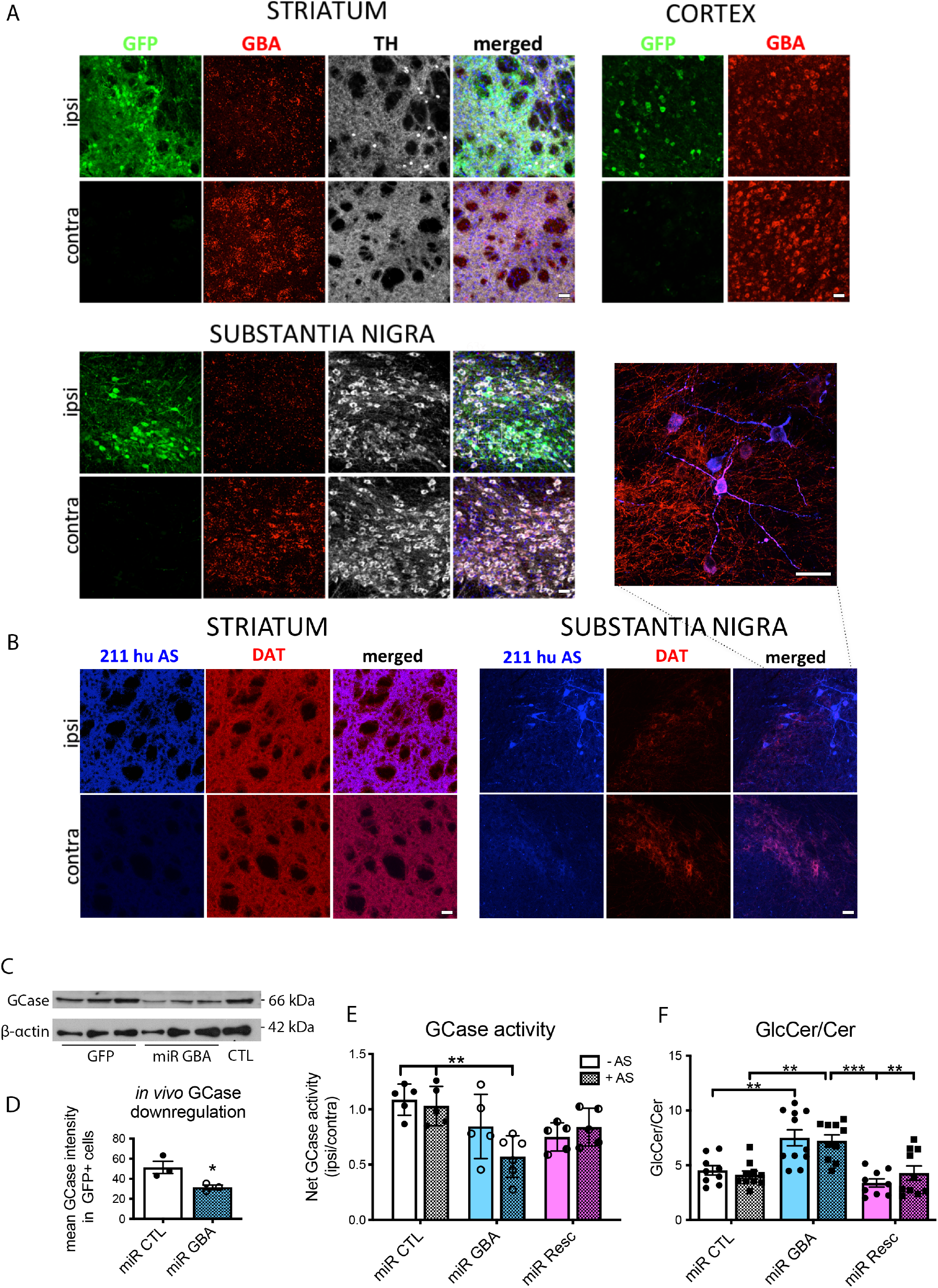
*In vivo* microRNA-mediated downregulation of GBA (miR GBA) +/- human α-synuclein (AS) in the striatum decreased GCase levels and activity and increased accumulation of glucosylceramide. Representative immunofluorescent images in the ipsi(lateral) (injected; top panel) vs. contra(lateral) (non-injected; bottom panel) hemisphere of 2 month-old wild-type mice 8 weeks post-injection (WPI) of microRNA (miR) GBA-GFP (green)-injected downregulation of GBA (red) in the striatum, cortex, and substantia nigra (clockwise); dopaminergic terminals (striatum) and cell bodies (substantia nigra) are indicated in pseudocolor with tyrosine hydroxylase (TH) antibody; scale bar: 25 μm; (A). Overexpression of human AS demonstrated with the 211 antibody (211 human (hu AS), blue) in dopamine transporter- (DAT; red) expressing striatal dopaminergic terminals (left) and cell bodies of the substantia nigra (right and top (merged image of nigral neuron at 63x magnification, 2x digital zoom); scale bar: 40 μm; (B). Western immunoblots demonstrating downregulation (~60%) of glucocerebrosidase (GCase) protein in Neuro2A cells overexpressing miR GBA vs. GFP and control (CTL) conditions with β-actin as a loading control (C); quantification). Quantification of *in vivo* downregulation (~40%) in miR GBA vs. miR CTL- injected mice cortices (mean GCase intensity (masked red) in GFP+ (green) cells) (p=0.038) (D). Measurement of GCase activity (net GCase activity ipsi/contra) revealed a statistically significant (treatment effect: p=0.0009) decrease in miR GBA + AS-injected mice vs. miR CTL +/- AS and miR rescue (miR Resc)-injected mice had similar levels to miR CTL, regardless of AS overexpression (E). The ratio of glucosylceramide to ceramide (GlcCer/Cer; ipsi/contra) was elevated in miR GBA-injected (treatment effect: p<0.0001) vs. miR CTL- and miR Resc-injected mice. *p<0.05, **p<0.001, ***p<0.0001. (F).

#### miRNA-mediated GCase downregulation and AS overexpression is accompanied by synergistic nigrostriatal neurodegeneration

Striatal dopamine levels (F(_2,25_)= 3.78, p=0.037, interaction: F(_2,25_)=3.32, p=0.053), TH immunostaining and TH+ neuron stereological counts in the substantia nigra (miR treatment: F(_2,34_)=11.08, p=0.0002) were decreased only in the miR GBA + AS group and both of these effects were reversed in the miR Resc + AS group (Fig 3A-C). The inverse relationship between GBA levels and TH expression (when combined with AS overexpression) in the substantia nigra is evident in immunofluorescent images (Fig 3D), indicating a specific synergistic neurodegenerative effect of GCase downregulation and AS overexpression.

**Figure 3.**
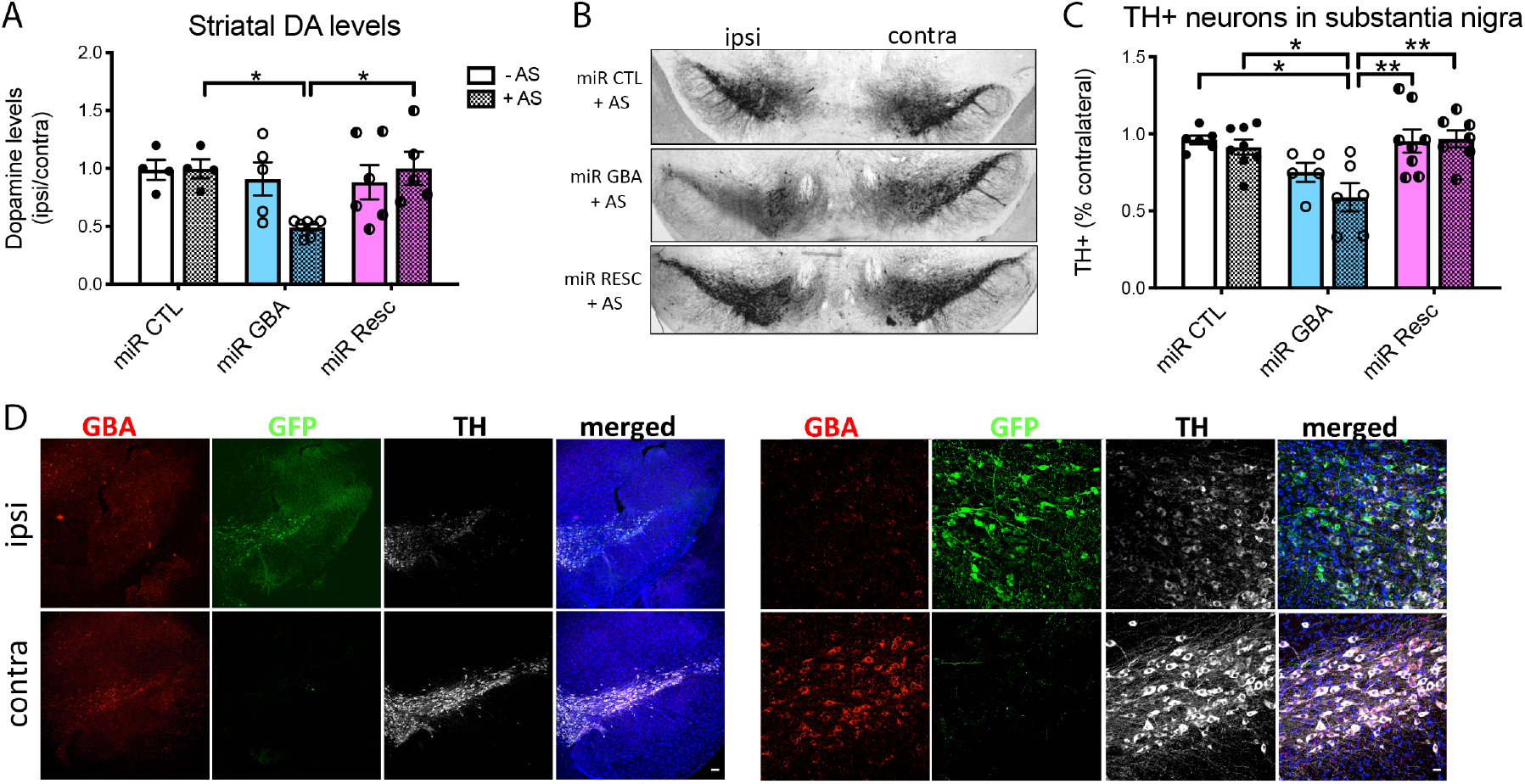
MicroRNA-mediated downregulation of GBA (miR GBA) in combination with human α-synuclein (AS) overexpression leads to dopaminergic nigrostriatal neurodegeneration. Unilateral injection of GFP-miR GBA + human AS (miR GBA + AS) in the striatum demonstrates: reduced striatal dopamine levels ((ipsi)lateral vs. (contra)lateral) compared with miR control + AS (miR CTL + AS) (p=0.047) and miR rescue + AS (miR Resc + AS) (p=0.027) (A); decreased tyrosine hydroxylase-positive (TH+) neurons in the substantia nigra on the (ipsi)lateral side of miR GBA + AS-injected mice as depicted in the representative images (10x) (B); quantification of stereological counts (% contra), demonstrates the same decrease compared with miR CTL +/- AS (p=0.021, 0.0061, respectively) and miR Resc +/- AS (p=0.0033, 0.0037, respectively) (C); and representative immunofluorescent images in the substantia nigra of the ipsi(lateral) (injected; top panel) vs. contra(lateral) (non-injected; bottom panel) hemisphere of miRNA GBA-GFP (green) and AS-injected 2 month old mice depicting downregulation of GBA (red) in TH+ cell bodies (pseudocolor) and merged images with DAPI nuclear staining (blue) (D); left panels 10x (scale bar: 100 μm), right panels 40x (scale bar: 25 μm); *p<0.05, **p<0.001.

#### Enhanced AS accumulation and release along the nigrostriatal axis mediated by GBA downregulation

Enhanced accumulation of human AS was evident in both the striatum and substantia nigra in the miR GBA + AS groups, while levels of AS in the miR Resc + AS group were comparable to miR CTL + AS levels (striatum: Fig 4A left, B; F(_2,12_)=6.54, p=0.012; substantia nigra: Fig 4A right, C; F(_2,23_)=26.22, p<0.0001). Next, human AS cDNA levels were measured in the ventral midbrain. The data revealed no differences in AS cDNA expression (Fig 4D), thus precluding differences in human AS levels due to the expression of the viral transgene in the various conditions. Finally, *in vivo* microdialysis in the striatum revealed enhanced AS release in the miR GBA + AS group compared to miR CTL + AS and this effect was reversed in the miR Resc + AS group (Fig 4E, F(_2,7_)=7.86, p=0.016). Increased AS expression in the miR GBA + AS group was accompanied by increased spontaneous activity measured in number of total rearings in the cylinder test (SF5, miR treatment: F(_2,59_)=10.09, p=0.0002, AS treatment: F(_1,59_)=4.17, p=0.046).

**Figure 4.**
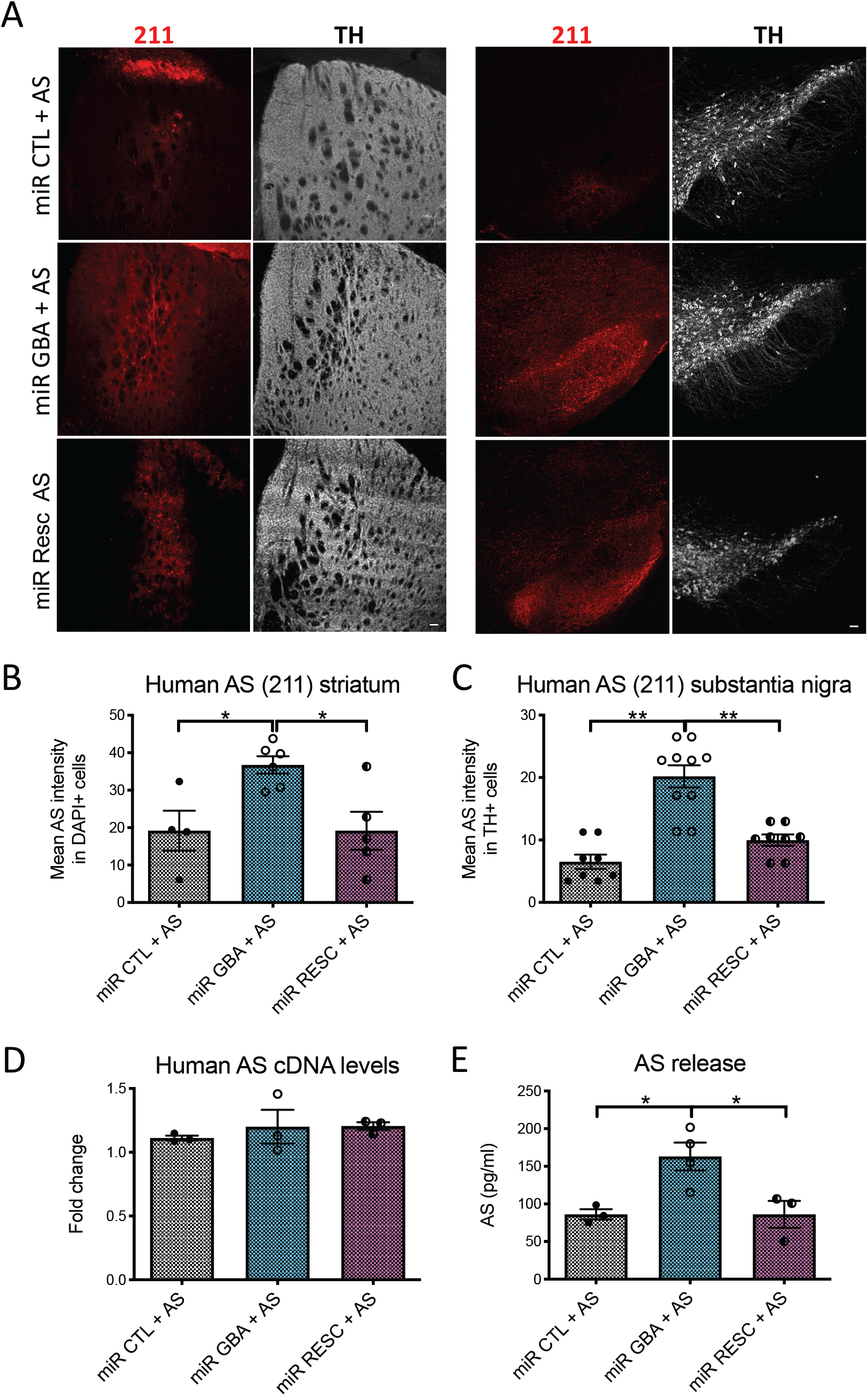
microRNA-mediated GBA (miR GBA) downregulation enhances human α-synuclein (AS) expression and release. Representative immunofluorescent images of human AS (211, red) staining in miR control (miR CTL)-, miR GBA- and miR rescue GBA (miR Resc)-injected in combination with human AS (+AS) in the striatum (left panel) and substantia nigra (right panel) with dopaminergic terminals and cell bodies, respectively, stained with tyrosine hydroxylase (TH, pseudo color) (A) (scale bar: 100 μm); quantification revealed a statistically significant increase in human AS expression following miR GBA + AS treatment in the striatum (p=0.012) (B); and in the substantia nigra (p<0.0001) (C) compared to miR CTL + AS and these effects were reversed in the miR Resc + AS group; cDNA levels of human AS, measured with qPCR (fold change) in ipsilateral vs. contralateral ventral midbrain tissue of the same mice, were not altered between the different treatment groups (D); in vivo microdialysis revealed enhanced extracellular AS release in the striatum of miR GBA + AS-injected mice compared to miR CTL + AS-injected mice and these effects were reversed in the miR Resc + AS group (E). *p<0.05, **p<0.001.

#### GCase downregulation is associated with astro- and microgliosis along the nigrostriatal axis

GFAP immunostaining in the striatum revealed increased astrogliosis following miR GBA treatment and this finding was significant in the miR GBA + AS group when compared with miR CTL + AS group and reversed with miR Resc expression (Fig 5A,B; miR treatment effect: F(_2,24_)=13.01, p=0.0001). GFAP immunostaining in the substantia nigra revealed a similar trend (Fig 5C,D; miR treatment effect: F(_2,24_)=3.95, p=0.033), with increased astrogliosis in the miR GBA-treated groups. Iba1 staining in the striatum revealed significantly increased microgliosis following miR GBA treatment, regardless of AS overexpression (Fig 5E,F; miR treatment effect: F(_2,23_)= 17.15, p<0.0001), while in the substantia nigra, microgliosis was enhanced only in the miR GBA-AS group and reversed in the miR Resc - AS group (Fig 5G,H; miR treatment effect: F(_2,24_)=6.54, p=0.0054). Overall, these data indicate that the main driver of inflammatory responses in this model, including both astrocytic and microglial activation, is the downregulation of GBA, while a synergistic effect of AS overexpression is perhaps slightly evident in the case of astrocytes.

**Figure 5.**
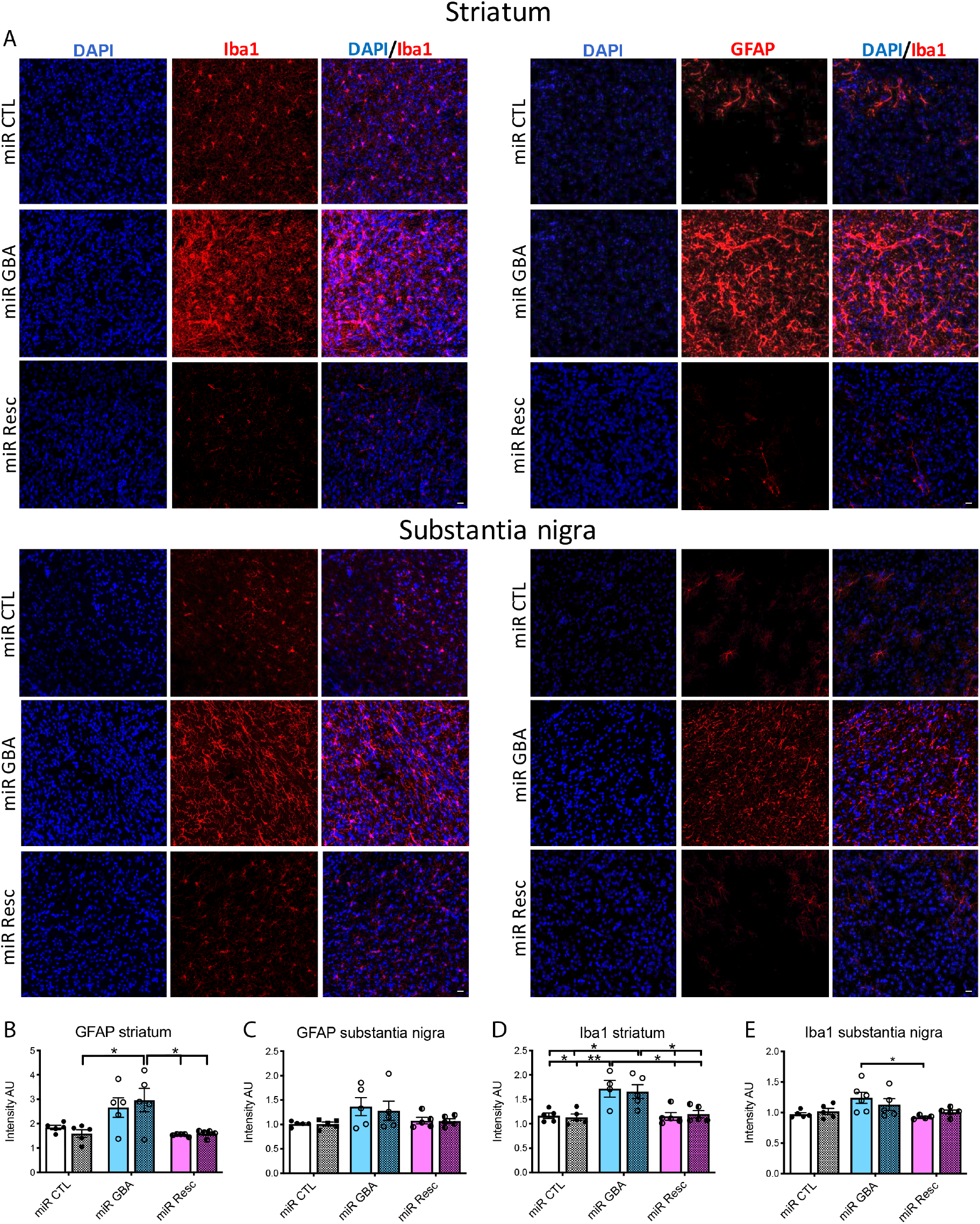
microRNA (miR)-mediated GBA (miR GBA) downregulation in the striatum induces astro- and microgliosis along the nigrostriatal axis. Representative immunofluorescent images of GFAP (marker for astrocytes, left panels) or AIF1/Iba1 (marker for microglia, right panels), DAPI (nuclear staining, blue) and merged images (DAPI/GFAP or DAPI/Iba1, respectively) in the striatum (upper panels) and substantia nigra (lower panels) of mice injected with miR control (miR CTL), miR GBA or miR rescue (miR Resc) (A) (scale bar: 25 μm). Quantification of GFAP intensity (expressed in arbitrary units (AU)) demonstrated increased expression in the mir GBA + α-synuclein (miR GBA + AS) group vs. miR CTL + AS and miR Resc +/- AS (p=0.0001) (B) in the striatum and the same trend persisted in the substantia nigra (p=0.033, no post hoc differences) (C); quantification of Iba1 intensity (expressed in AU) revealed increased expression in miR GBA +/- AS compared to miR CTL +/- AS groups and effective reversal in miR Resc +/- AS groups (p<0.0001) in the striatum (D); in the substantia nigra, Iba1 expression was increased in the miR GBA – AS group vs. miR Resc – AS group (p=0.0054) (E). *p<0.05, **p<0.001.

## Discussion

In an effort to develop a novel mouse model which recapitulates GBA-PD pathogenesis, where *GBA* mutations are a risk factor but not a determinant of the disease process, our study uses viral models *in vivo* to assess the isolated and combined effects of modulating GBA activity and AS expression along the nigrostriatal axis. We demonstrate that neither WT nor N370S GCase overexpression alters endogenous AS levels, but that GCase downregulation via a microRNA results in insufficient GCase function, substrate accumulation, and neuroinflammation (astro- and microgliosi). Furthermore, the combination of GCase downregulation and human AS overexpression leads to a synergistic augmentation of AS burden and increased striatal AS release, resulting in significant nigrostriatal neurodegeneration. This double-hit model recapitulates key pathological features of PD which manifest at 8 WPI in young adult WT mice, rendering it a valuable tool for studying the underlying mechanisms of disease pathogenesis and screening of potential disease-modifying therapeutics.

Our assessment of WT and N370S GBA overexpression revealed minimal effects on AS tissue levels in the cortex and no effect on nigrostriatal dopaminergic system integrity. Mutant N370S GBA decreased secreted AS *in vitro*, perhaps indicating a disruption of the secretory pathway through effects of the mutant protein on the ER/Golgi/endosomal/lysosomal compartment. On the other hand, *in vivo* mutant N370S GBA, but not WT GBA viral expression, in the striatum of transgenic A53T AS mice had the opposite effects, enhancing extracellular AS release (Papadopoulos et al., 2018), suggesting that *in vivo* and cell culture-based mechanisms of AS secretion may be different. Of note, N370S heterozygosity did not affect the disease course in A30P AS mice (Taguchi et al., 2017). Thus, although *GBA* mutations alone may disrupt AS proteostasis, depending on the context, an enhanced AS burden is required to cause nigrostriatal neurodegeneration.

In our next set of experiments, we focused on the combination of miRNA-mediated GBA downregulation and human AS overexpression (miR GBA + AS). In this model, we observed a lack of significant GBA downregulation at 4 weeks, while at 8 weeks, striatal GCase was reduced by approximately 40% (Fig.2D), similar to levels present in GBA heterozygosity patients and models (Ikuno et al., 2019; Osellame et al., 2013). Furthermore, reduced GCase expression resulted in decreased GCase enzyme activity, more evident in the miR GBA + AS group, although GlcCer levels were elevated in miR GBA-treated mice, regardless of AS expression. Interestingly, the miR GBA + AS group exhibited elevated levels of human AS both in the striatum and substantia nigra and enhanced striatal AS release when compared with miR CTL + AS and miR Resc+ AS groups, further supporting the hypothesis of a “toxic” synergistic interplay. Both the miR CTL + AS and miR Resc + AS groups appeared to have similar levels of human AS expression, demonstrating that reversal of the phenotype and restoration of normal GCase function by overexpression of the human *GBA* gene is sufficient to prevent the aberrant accumulation of AS. We confirmed that this enhanced accumulation of AS was due to the downregulation of GBA along the nigrostriatal axis and not due to variance of the viral load delivered in the striatum of the animals, as all groups had similar levels of human AS mRNA. From our findings, it is apparent that when there is a reduction in the GCase activity of neurons overexpressing AS, there is enhanced AS accumulation. AS accumulation could be facilitated by GlcCer accumulation, due to deficits in GCase activity (Ikuno et al., 2019; Osellame et al., 2013). Neuronal cultures, derived from the knock-in L444P GBA mutant mouse expressing human AS, resulted in decreased human AS degradation and subsequently, increased AS steady state levels (Fishbein et al., 2014). Finally, deficits in lysosomal function and thus AS clearance may also play a role (Gegg et al., 2012; Schöndorf et al., 2018), leading to an overall increase of AS’s half-life. Thus, baseline AS burden is a prerequisite for a “snowball” effect to occur in the presence of lipid substrate accumulation and vice versa, reduced GCase expression/activity exerts gene dose-dependent effects on AS accumulation. It is of note that our findings here in this regard are at odds with our data in cultured neuronal SH-SY5Y cells and primary cortical neurons, where near-complete pharmacological inhibition of GCase activity did not alter the burden of AS, be it monomeric or oligomeric, endogenous or overexpressed (Dermentzaki et al., 2013). This may reflect, as above, differences between the cell culture and *in vivo* settings, but also may relate to the timing of exposure to these manipulations, which is much more prolonged in the present experiments, and the degree of accumulation of the glycolipid substrates in each case.

Heterozygous *GBA1* KO mice crossed with AS BAC mice also demonstrated reduced GCase activity, however, GlcCer levels and total AS levels were unaffected while mild nigral dopaminergic cell loss and phosphorylated AS were only observed in aged (>16 month) mice (Ikuno et al., 2019). Moreover, heterozygous GBA1 KO mice crossed with A53T AS mice had neither a difference in GCase levels and activity nor in AS levels, even though symptom severity and lethality were accelerated (Tayebi et al., 2017). Taken together, unknown indirect mechanisms of GBA loss-of-function and potential developmental compensatory mechanisms in transgenic mice exist that may not affect viral-mediated altered expression in adult mice. Along these lines, GCase deficiency caused by L444P heterozygosity is not itself sufficient to induce nigral cell loss, but it significantly enhances the toxic effects of AS overexpression (Ikuno et al., 2019; Osellame et al., 2013). The L444P mutation prolongs the half-life of endogenous AS and AAV-delivered human AS, thus enhancing its intraneuronal levels and promoting AS assembly (Migdalska-Richards et al., 2017).

Post-mortem GBA-PD brains demonstrate extensive neuroinflammation and significant loss of dopaminergic nigral neurons (Wang et al., 2015). In murine models of GBA deficiency (both transgenic and pharmacologically induced with CBE), microglial activation and astrogliosis are spatially and temporally correlated with AS accumulation (Ginns et al., 2014; Kim et al., 2018; Yun et al., 2018) and nigrostriatal degeneration when there is a PD challenge such as MPTP (Yun et al., 2018) or AS overexpression (A53T mice) (Kim et al., 2018). Our findings support a link between GBA deficiency and enhanced glial activation and astrogliosis along the nigrostriatal axis (Fig. 4). However, excess AS accumulation and enhanced striatal release does not appear to further exacerbate this inflammatory response; rather, the synergistic effects of GBA deficiency and AS overexpression is required for full-blown nigrostriatal degeneration. Based on previous studies suggesting that extracellular AS can trigger an inflammatory response by increasing proinflammatory cytokines and glial activation (Chung et al., 2009; Engelender and Isacson, 2017; Theodore et al., 2008), our results suggest that a ceiling effect may be achieved with GBA deficiency on nigrostriatal neuroinflammation. However, GBA deficiency in the presence of excess AS drives its further accumulation and deposition into the extracellular space, potentially contributing to the neurodegenerative process.

GBA-PD correlates with more severe motor phenotypes and increased risk of nonmotor manifestations, including dementia and psychosis (Gegg et al., 2012; Schöndorf et al., 2018). Interestingly, assessment of spontaneous motor activity revealed a synergistic enhancement of locomotor activity with GBA downregulation and human AS overexpression, reminiscent of psychosis, as we have previously reported in human AS BAC transgenic rats (Polissidis et al., 2020). Similar findings were observed in AS BAC transgenic mice crossed with *GBA* heterozygous mice (Ikuno et al., 2019). Interestingly, no unilateral impairments were observed, but it is possible that the synergistic effects of GBA downregulation and AS overexpression, on excess AS accumulation and striatal release, over-ride potential subtle motor impairments.

## Conclusions

The present data support a loss-of-function association of *GBA* mutations in PD and suggest that GBA-PD is more likely to develop if AS burden is increased. Such an increase could concomitantly occur through genetic or epigenetic regulation, and could be further driven by GBA partial deficiency, as demonstrated here, leading to the PD phenotype manifestation, once a threshold is crossed. Such models are vital to study the pathogenic mechanisms involved in GBA-associated nigrostriatal neurodegeneration in PD. Most importantly, understanding such mechanisms may lead to novel therapeutic targets for GBA-PD, but also potentially, more broadly in sporadic PD.

This novel PD model has several advantages over current models including construct validity (genetics and pathology) and the manifestation of a neurodegenerative phenotype within a short time frame (2 months); ideal for assessing both potential preventative and disease-modifying therapeutic strategies. Therapeutic development efforts are currently focused on boosting GCase activity, reducing substrate accumulation or aberrant AS. The propensity of GCase deficiency to lower the threshold for pathological AS accumulation and subsequently, promote its release into the extracellular space, thus spreading pathology, supports the therapeutic potential of AS immunotherapy. On the other hand, in a recent phase 2 study, venglustat, a glucosylceramide synthase inhibitor aimed at reducing glycolipid substrate accumulation, failed to demonstrate efficacy in improving the Unified Parkinson’s Disease Rating Scale score in GBA-PD patients. This major therapeutic disappointment, however, demonstrates GBA’s complex role in PD beyond substrate accumulation and highlights the importance and urgency of deciphering novel AS-GBA-based pathway interactions at the molecular, cellular, and systems level in robust models of GBA-linked PD.

## Acknowledgments

We would like to acknowledge the technical support of the Animal Housing facility personnel and Dr. Stamatis Pagakis and Eleni Rigana of the Bioimaging Unit at BRFAA.

## Author Contributions

Alexia Polissidis: Conceptualization, Data curation, Formal analysis, Funding acquisition, Investigation, Methodology, Resources, Supervision, Writing-Original draft preparation.: Georgia Nikolopoulou: Conceptualization; Formal analysis; Investigation; Methodology; Writing-Original draft preparation.: Effrosyni Koronaiou: Formal analysis; Investigation; Visualization.: Maria Nikatou: Formal analysis; Investigation.: Catherine Viel: Formal analysis; Investigation.: Modestos Nakos-Bimpos: Formal analysis; Investigation; Visualization.: Marios Boyiongko: Formal analysis; Investigation, Supervision: S. Pablo Sardi: Resources, Supervision, Validation; Methodology; Writing - review & editing.: Konstantinos Vekrellis: Funding acquisition; Resources; Writing - review & editing.: Maria Xilouri: Conceptualization; Methodology; Writing - review & editing.: Leonidas Stefanis: Conceptualization; Funding acquisition; Methodology; Project administration; Resources; Supervision; Writing - review & editing.

