## Supplementary Files for "A double-hit *in vivo* model of *GBA1* viral microRNA-mediated downregulation and human alpha-synuclein overexpression demonstrates nigrostriatal degeneration"

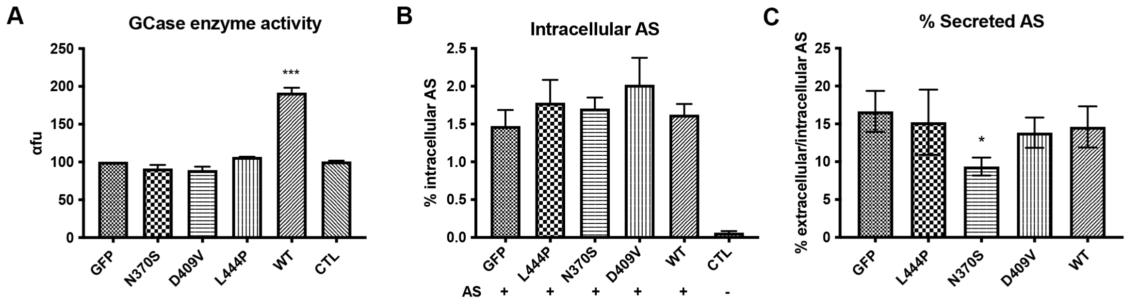


SF1. *In vitro* viral overexpression of GFP, wild-type (WT) and mutant (N370S, D409V, L444P) GBA. WT GBA, but not mutant overexpressing HEK293 cells, led to enhanced glucocerebrosidase (GCase) enzyme activity when compared to control (CTL)-treated cells (expressed in arbitrary fluorescence units (afu) ($p<0.0001$) (A); no changes were observed in levels of intracellular α-synuclein (AS) (expressed as % intracellular levels vs. total AS levels) ($p=0.617$) (B); N370S mutant GBA decreased % secreted AS (expressed as % extracellular AS levels vs. intracellular AS levels) ($p=0.037$) (C); *$p<0.05$, ***$p<0.0001$.

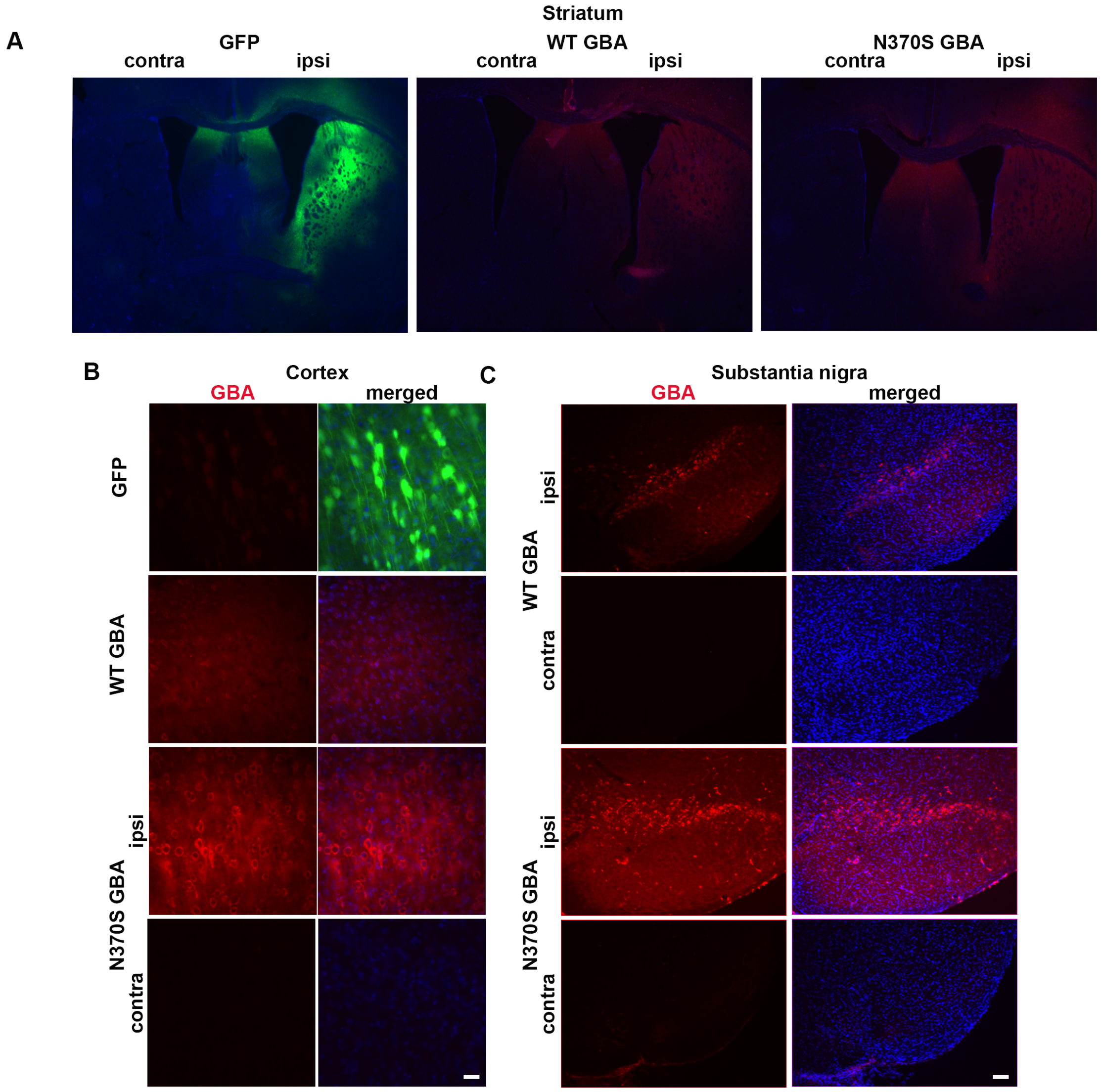


SF2. Viral gene expression in the striatum, cortex, and substantia nigra following unilateral striatal injections of GFP, wild-type GBA (WT GBA) or N370S GBA adeno-associated viruses (AAVs in 2-month old WT mice 8 weeks post-injection. Representative immunofluorescent images in the ipsi(lateral) (injected; right) and contra(lateral) (non-injected; left) hemisphere of GFP (green), wild-type (WT) GBA, and N370S GBA (red) AAV delivery in the striatum (2.5x magnification); merged images include DAPI nuclear staining (blue) (A); expression in the cortex (scale bar: 25 µm), and retrograde transport to the substantia nigra (scale bar: 50 µm) (C).

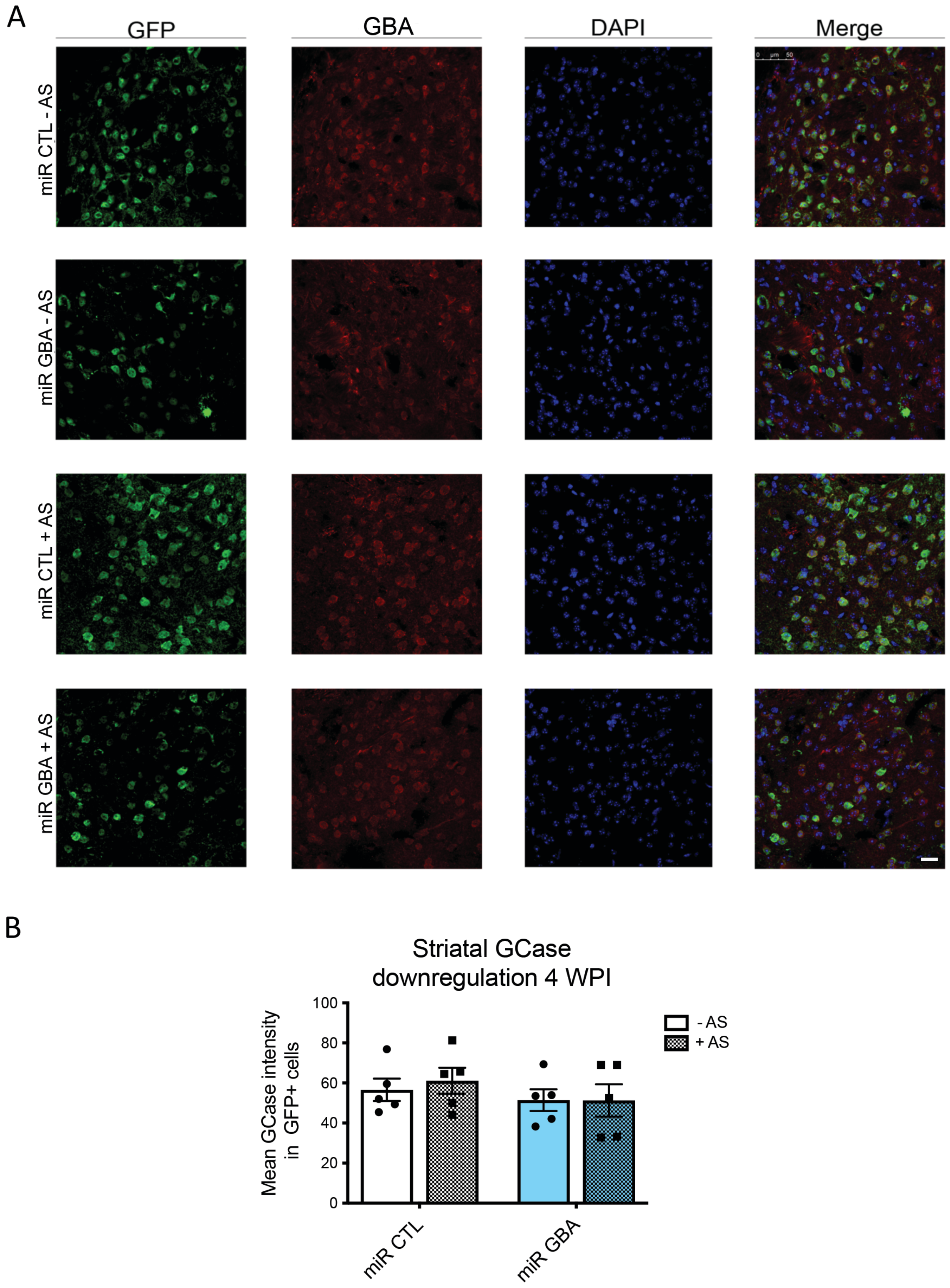

A
GFP
GBA
DAPI
Merge
miR CTL - AS
miR GBA - AS
miR CTL + AS
miR GBA + AS
B
Striatal GCase downregulation 4 WPI
Mean GCase intensity in GFP+ cells
100
80
60
40
20
0
- AS
+ AS
miR CTL
miR GBA

SF3. *In vivo* microRNA-mediated downregulation of GBA (miR GBA) +/- human α-synuclein (AS) in the striatum did not affect GCase expression at 4 weeks post-injection (WPI). Representative immunofluorescent images in the ipsilateral hemisphere of 2 month-old wild-type mice striata injected with either miR GBA-GFP (miR GBA) (green) or miR control (CTL)- GFP (miR CTL) (green) +/- AS and expression of GBA (red) in the striatum, including DAPI nuclear staining (blue) (scale bar: 40 µm) (A). Quantification of mean GCase (GBA) expression (intensity in GFP+ cells) at 4 WPI revealed a nonsignificant incremental decrease in GCase downregulation.

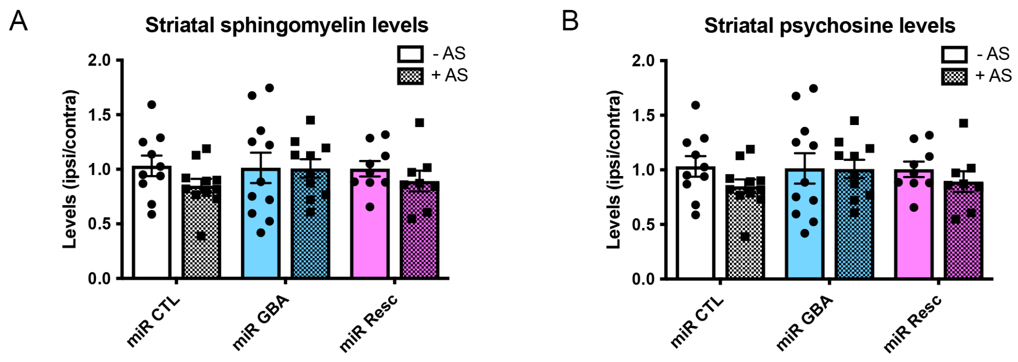


SF4. Glycosphingolipid levels in the striatum of 2-month old wild-type mice injected unilaterally with microRNA control (miR CTL), GBA (miR GBA) or rescue (miR Resc) in combination with human α-synuclein (AS). Both sphingomyelin (A) and psychosine (B) levels were unaffected by the experimental treatments (levels expressed as µg/g (ipsi)lateral/(contra)lateral striatum).

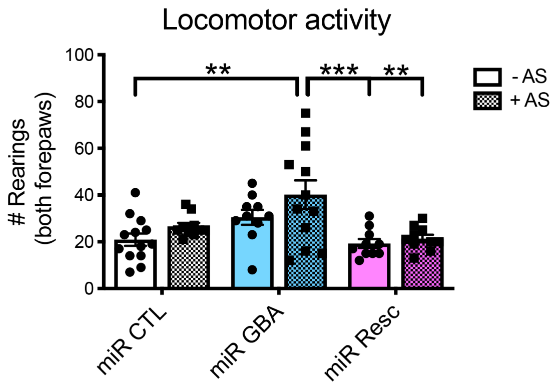


SF5. Locomotor activity of 2-month old wild-type mice injected unilaterally with microRNA control (miR CTL), GBA (miR GBA), or rescue (miR Resc) in combination with human α-synuclein (AS) in the striatum. The number of rearings in a 5-min cylinder test were increased following miR GBA + AS treatment as compared to miR CTL – AS ($p=0.0012$), miR Resc – AS ($p=0.001$), and miR Resc + AS ($p=0.0039$). **$p<0.01$, ***$p<0.001$.
